# Loss of Propionyl-CoA Carboxylase Reprograms Hepatic Metabolism by Suppressing Mitochondrial Pyruvate Carboxylation and Fatty Acid Oxidation

**DOI:** 10.64898/2026.04.13.718201

**Authors:** Fang Lu, Chorlada Paiboonrungruang, Wentao He, Zhaohui Xiong, Pingchuan Tang, Takhar Kasumov, Xiaoxin Chen, Guo-Fang Zhang

## Abstract

Propionic acidemia (PA) is an inborn error of metabolism caused by propionyl-CoA carboxylase (PCC) deficiency due to mutations in either *PCCA* or *PCCB*. Without proper management, the disease is associated with high mortality. Even with dietary restriction, patients often develop complications later in life, and the underlying pathological mechanisms remain poorly understood. The liver is the primary organ responsible for propionyl-CoA metabolism, yet the metabolic alterations induced by PCC deficiency in the liver have not been systematically investigated. In this study, we used a hepatocyte model of PA— *PCCA^null^-*HepG2 cells—to comprehensively examine metabolic alterations using stable isotope–based metabolic flux analysis. The *PCCA* knockout recapitulated key metabolic features of PA in HepG2 cells. Furthermore, *PCCA* deficiency reduced mitochondrial fatty acid oxidation while increasing glucose oxidation through pyruvate dehydrogenase. In contrast, pyruvate anaplerosis via pyruvate carboxylase was markedly reduced in *PCCA* knockout cells. This reduction in anaplerotic flux impaired the capacity for gluconeogenesis and lipid synthesis, consistent with observations from *in vivo* studies in *Pcca⁻/⁻* (A138T) mice. Additionally, branched-chain keto acid catabolism was reduced in *PCCA* knockout HepG2 cells. Threonine showed minimal metabolic contribution in this model, further supporting the role of propionate as a major source of propionyl-CoA production. Collectively, these findings highlight the metabolic vulnerabilities associated with PCC deficiency and underscore the increased risk of prolonged fasting in patients with PA, particularly those with severe disease.

## Introduction

Propionic acidemia (PA) is a severe inborn error of metabolism caused by loss-of-function mutations in *PCCA* or *PCCB*, which encode the two subunits of propionyl-CoA carboxylase (PCC) ^1–16^. The global incidence of PA ranges from approximately 1 in 50,000 to 1 in 100,000 live births, with an estimated prevalence of ∼1 in 100,000 individuals (∼3,400 people) in the United States ^8,12^. Most cases are diagnosed within the first three months of life, and delayed diagnosis is associated with severe developmental delay and multi-organ complications. Despite medical management, mortality remains high (7–12%) ^12^.

PA is characterized by a broad spectrum of complications, including cardiac dysfunction, neurological impairment, renal disease, immune abnormalities, pancreatitis, and liver dysfunction ^8^. However, the organ-specific metabolic mechanisms underlying these complications remain incompletely understood. At the biochemical level, PCC deficiency leads to accumulation of propionyl-CoA, which disrupts metabolic homeostasis by depleting free CoA and carnitine pools—both essential for nutrient oxidation and energy production ^17–19^. In addition, propionyl-CoA competes with acetyl-CoA in multiple metabolic pathways. Under physiological conditions, acetyl-CoA levels exceed propionyl-CoA by more than tenfold, minimizing this competition; however, in PA, elevated propionyl-CoA reduces acetyl-CoA availability and generates toxic metabolites such as methylcitrate and propionylglutamate ^20^. These metabolites impair ammonia detoxification and disrupt the tricarboxylic acid (TCA) cycle.

Importantly, propionyl-CoA metabolism is highly organ-dependent. Organs such as liver, kidney, and pancreas exhibit high propionyl-CoA metabolic capacity, whereas skeletal muscle has relatively limited activity ^21,22^. Therefore, understanding PA pathophysiology requires investigation in organ-specific models.

Although mouse models have provided valuable insights, their translational limitations are well recognized ^23,24^. Human cell-based systems, including cell lines and organoids, offer cost-effective and clinically relevant alternatives. Our previous work demonstrated that *PCCA* deficiency induces a metabolic shift from fatty acid oxidation to glucose utilization in hiPSC-derived cardiomyocytes, a phenotype consistent with failing hearts^17^.

The liver plays a central role in systemic metabolism and is the primary site of propionyl-CoA clearance ^25^. It is also a key target for transplantation in severe PA ^7,26–32^. However, the metabolic consequences of PCC deficiency in human hepatocytes remain poorly defined, since Chapman *et al.* created a first successful culture of PA patient-derived primary hepatocytes for the purpose of better understanding of the molecular pathophysiology of PA and examining the effectiveness of potential therapeutic agents in the most relevant tissue ^33^.

To address this gap, we generated a CRISPR-mediated *PCCA* knockout in the HepG2 cells (*PCCA^null^*-HepG2) and performed stable isotope–based metabolic flux analyses. Our data demonstrate that *PCCA* deficiency reduces pyruvate carboxylation while increasing pyruvate oxidation via pyruvate dehydrogenase (PDH), leading to impaired anaplerosis, reduced gluconeogenesis and fatty acid synthesis capacity, and altered substrate utilization. These findings provide mechanistic insight into liver dysfunction in PA and identify potential metabolic targets for therapeutic intervention.

## Results

### *PCCA* knockout recapitulates the metabolic phenotype of PA

A CRISPR guide RNA was used to knock out PCCA and generate PCCA-null HepG2 cells as a hepatic model of PA (Supplementary Fig. 1). CRISPR-mediated knockout of *PCCA* resulted in undetectable PCCA protein (Fig. 1A) and PCC enzymatic activity in HepG2 cells compared with wild-type (WT) controls (Fig. 1B). Consistent with this loss of function, *PCCA^null^*-HepG2 cells exhibited marked accumulation of propionyl-CoA and propionylcarnitine, along with an increased C3/C2 acyl-CoA ratio and elevated methylcitrate levels (Figs. 1C–1F). These metabolic alterations are characteristic of PA and confirm that the cellular model reproduces key biochemical features of the disease ^8,21^. This model therefore provides a suitable platform to investigate liver-specific metabolic consequences of PCC deficiency.

**Figure 1.**
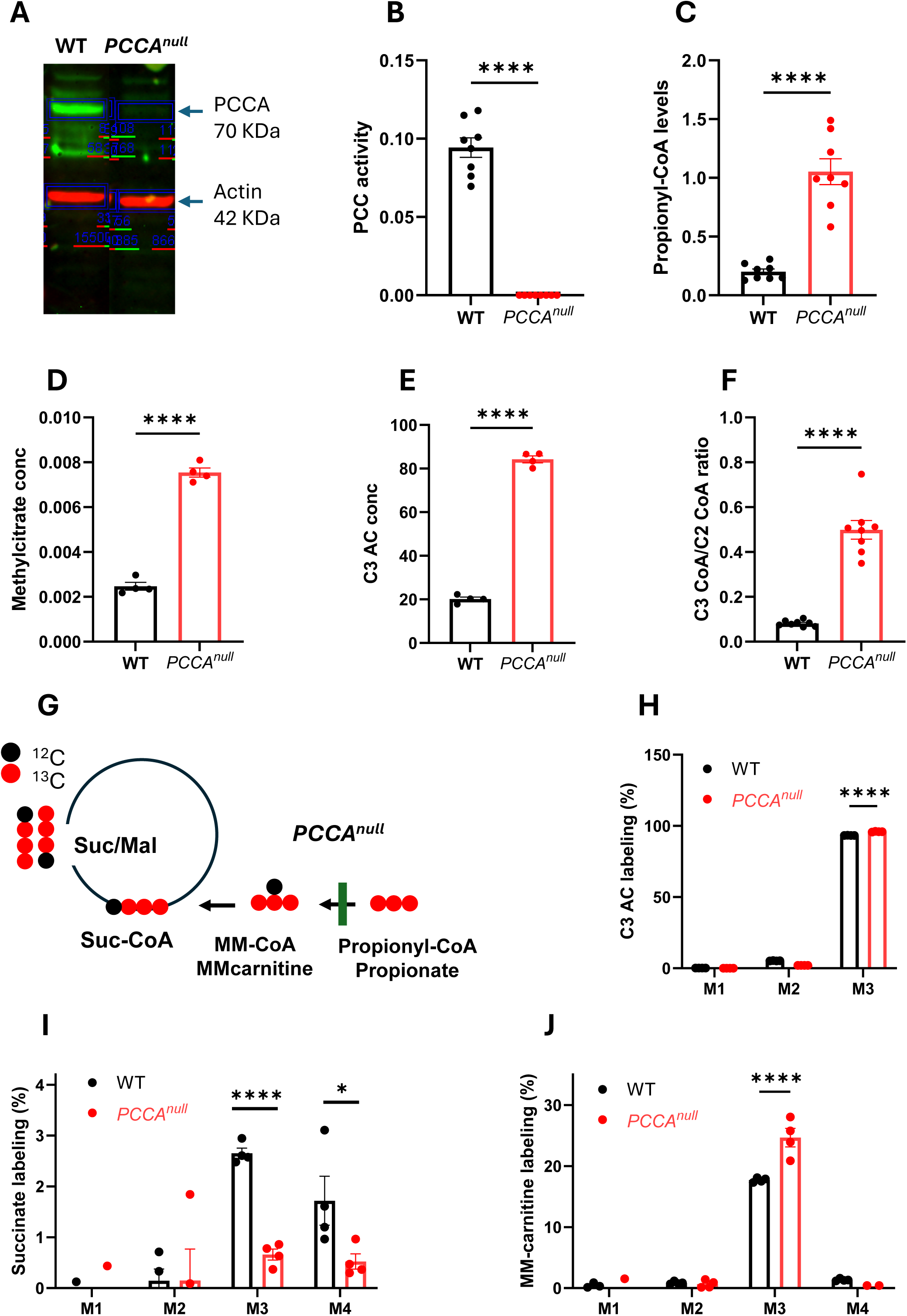
*PCCA^null^*-HepG2 cells recapitulate key metabolic features of propionic acidemia. (A) Western blot analysis of *PCCA* in WT and *PCCA* knockout (KO) HepG2 cells. (B) Propionyl-CoA carboxylase (PCC) activity in WT and *PCCA^null^*-HepG2 cells. (C–E) Levels of propionyl-CoA, methylcitrate, and C3 acylcarnitine (C3 AC) in WT and *PCCA^null^*-HepG2 cells, respectively. (F) Ratio of propionyl-CoA to acetyl-CoA (C3-CoA/C2-CoA) in WT and *PCCA^null^*-HepG2 cells. (G) Schematic of [¹³C₃]propionate tracing into the TCA cycle. (H–J) Isotopomer labeling of propionylcarnitine, succinate, and methylmalonylcarnitine (MM-carnitine) in WT and *PCCA^null^*-HepG2 cells following incubation with 1 mM [¹³C₃]propionate for 4 hours. Data are presented as mean ± SEM (n = 4). *, **, and **** indicate p < 0.05, p < 0.01, and p < 0.0001, respectively.

### Impaired propionyl-CoA anaplerosis revealed by [¹³C₃]propionate tracing

To investigate propionyl-CoA metabolism, WT and *PCCA^null^*-HepG2 cells were incubated with 1 mM [¹³C₃]propionate for 4 hours, followed by analysis of downstream metabolite labeling (Fig. 1G). As expected, propionylcarnitine was predominantly labeled as the M3 isotopomer (>93%) in both groups, with slightly higher enrichment in *PCCA^null^*-HepG2 cells, consistent with impaired downstream metabolism (Fig. 1H).

In contrast to *in vivo* liver metabolism ^21^, incorporation of propionate-derived carbon into TCA cycle intermediates was limited in HepG2 cells, as reflected by low labeling (∼2–3%) of succinate, compared to ∼20% observed *in vivo* ^21^ (Fig. 1I). Notably, *PCCA^null^*-HepG2 cells exhibited a marked reduction in labeling of all TCA cycle intermediates (Supplementary Figs.2A-2D), indicating impaired anaplerotic entry of propionyl-CoA into the TCA cycle.

Despite complete loss of PCC activity, low levels of M3 labeling were still detected in downstream metabolites (Fig. 1I). This observation was further supported by the presence of M3 methylmalonylcarnitine derived from [¹³C₃]propionate (Fig. 1J and Supplementary Fig. 2E). The isotopic labeling pattern and corresponding mass spectra of methylmalonylcarnitine under labeled propionate and unlabeled conditions are shown Supplementary Figs. 2F and 2G). Together, these data suggest that a small fraction of propionyl-CoA may still be converted to methylmalonyl-CoA through alternative enzymatic activity. Although this flux is minimal, it points to the potential involvement of other acyl-CoA carboxylases, which warrants further investigation.

### *PCCA* deficiency shifts pyruvate metabolism toward PDH

To determine how PCC deficiency affects glucose metabolism, we performed [¹³C₆]glucose tracing in WT and *PCCA^null^*-HepG2 cells (Fig. 2A). Labeling of glycolytic intermediates, including pyruvate, lactate, and alanine, was slightly decreased or unchanged in *PCCA^null^*-HepG2 cells (Figs. 2B–2D), indicating that glycolytic flux was largely unaltered. This was further supported by other glycolic intermediates labeling or levels (Supplementary Figs. 3A-F). In contrast, M2 acetylcarnitine labeling and levels was significantly increased without changing endogenous unlabeled levels of acetylcarnitine (Fig. 2E and Supplementary Figs. 3G-3H), suggesting enhanced conversion of pyruvate to acetyl-CoA via PDH. This increase in PDH flux was further supported by elevated M2 citrate labeling (Fig. 2F). However, higher-order citrate isotopomers (M3, M5, and M6) were reduced or showed a downward trend (Fig. 2F), resulting in no significant change in total citrate labeling (Fig. 2G).

**Figure 2.**
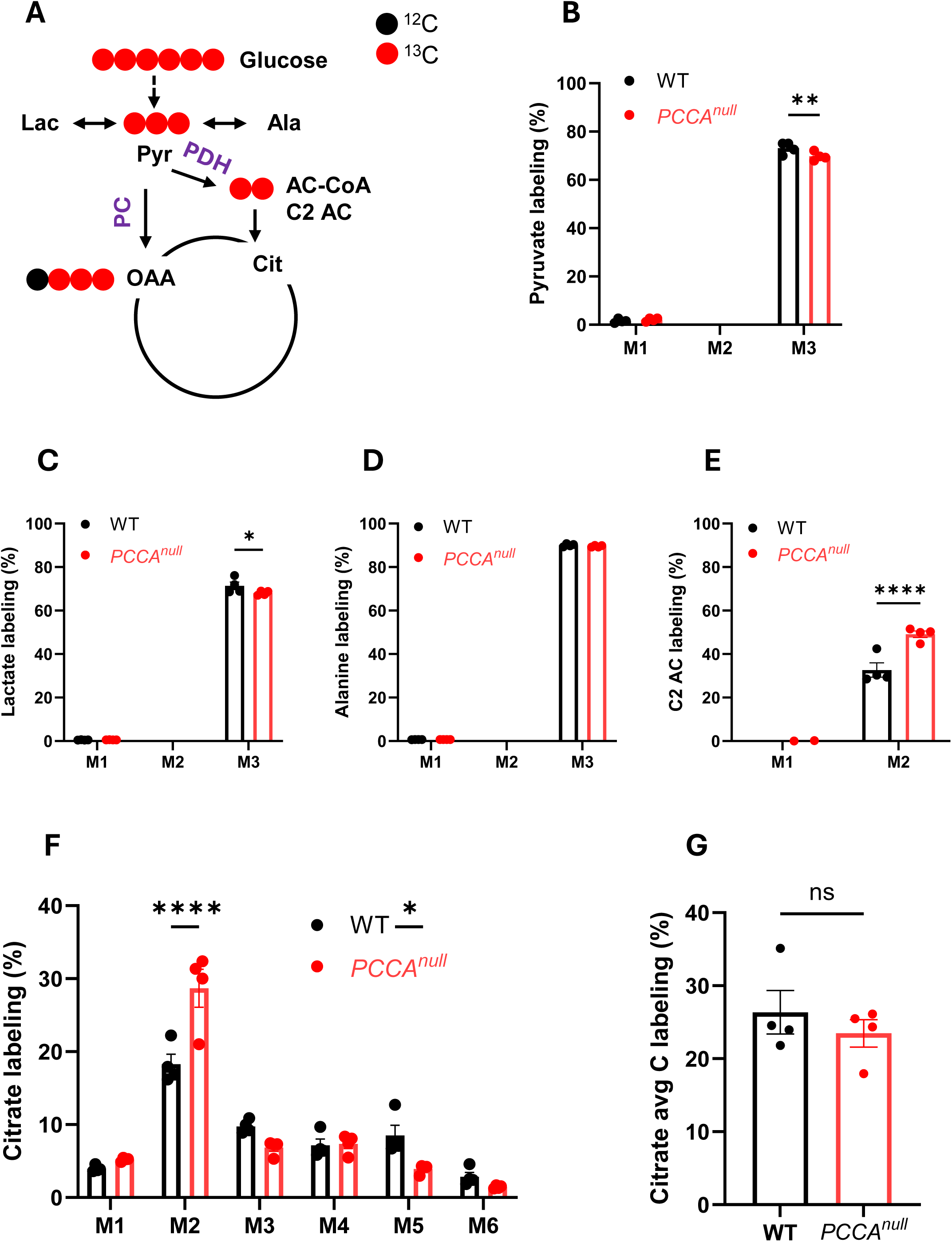
*PCCA* knockout enhances glucose metabolism through pyruvate dehydrogenase. (A) Schematic of [¹³C₆]glucose metabolism entering the TCA cycle via pyruvate dehydrogenase (PDH) and pyruvate carboxylase (PC). (B–F) Isotopomer labeling of pyruvate, lactate, alanine, acetylcarnitine (C2 AC), and citrate in WT and *PCCA^null^*-HepG2 cells. (G) Average carbon (Avg C) labeling of citrate. Data are presented as mean ± SEM (n = 4). *, ***, and **** indicate p < 0.05, p < 0.005, and p < 0.0001, respectively.

The discordant changes between M2 citrate and higher-order isotopomers suggest additional alterations in TCA cycle flux beyond acetyl-CoA entry from pyruvate. Further analysis of TCA cycle intermediates showed that M3 isotopomers of malate, aspartate, and fumarate predominated in WT cells but were markedly reduced in *PCCA^null^*-HepG2 cells (Fig. 3A and Supplementary Figs. 4A–4C), consistent with active pyruvate carboxylation mediated by pyruvate carboxylase (PC) in hepatic cells. Notably, PCCA deficiency impairs pyruvate anaplerosis via PC.

**Figure 3.**
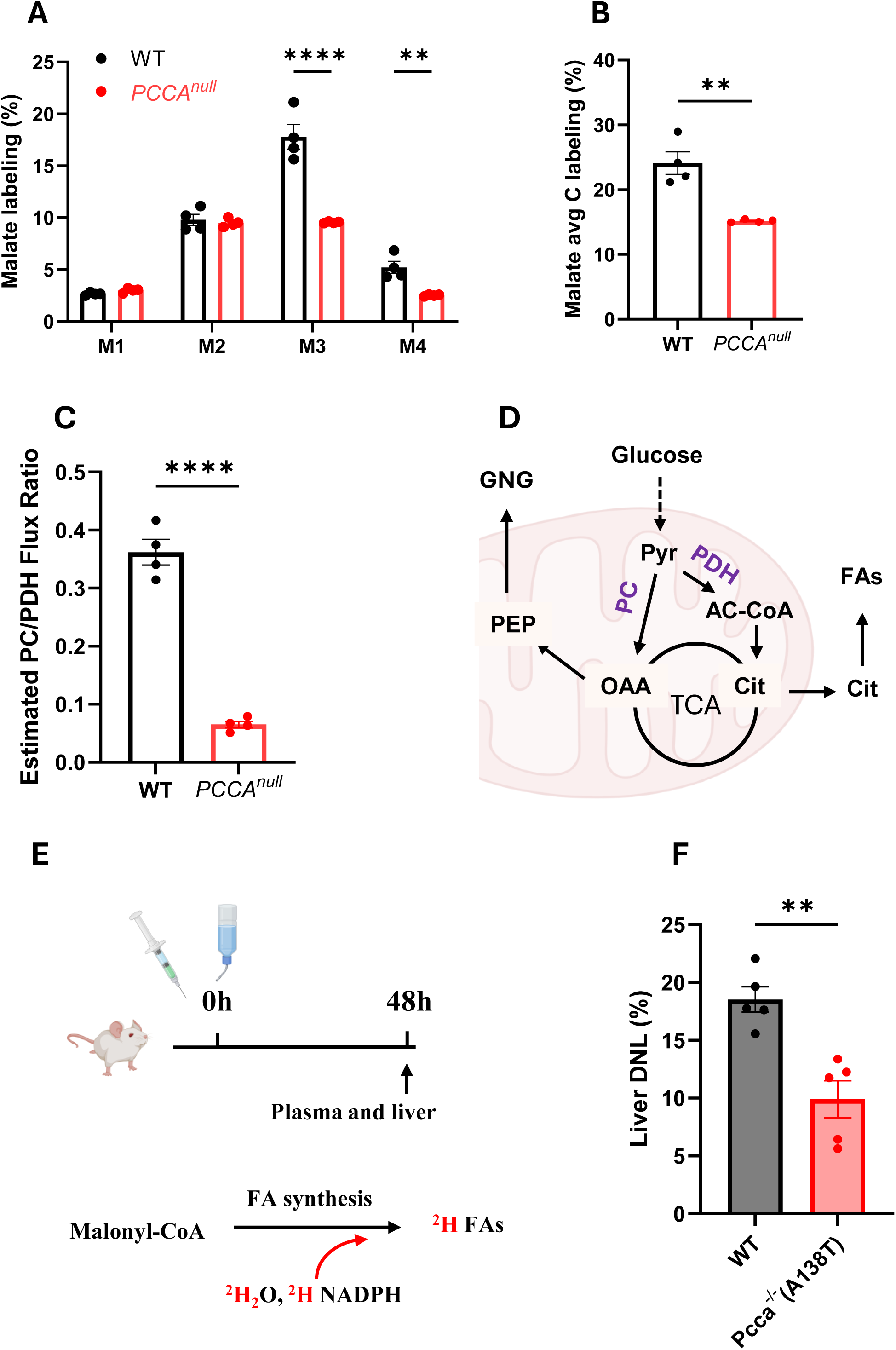
*PCCA* knockout markedly reduces pyruvate carboxylation and de novo lipogenesis. (A–B) Mass isotopomer distributions of malate and aspartate in WT and *PCCA^null^*-HepG2 cells following incubation with 11 mM [¹³C₆]glucose for 4 hours. (C) Estimated relative flux ratio of pyruvate carboxylase (PC) to pyruvate dehydrogenase (PDH). (D) Schematic of glucose metabolism supporting fatty acid synthesis and gluconeogenesis (GNG). (E) Experimental design for measuring DNLin mice. (F) Quantification of DNL. Data are presented as mean ± SEM (n = 4 for A–C; n = 5 for F). *, ***, and **** indicate p < 0.05, p < 0.005, and p < 0.0001, respectively.

**Figure 4.**
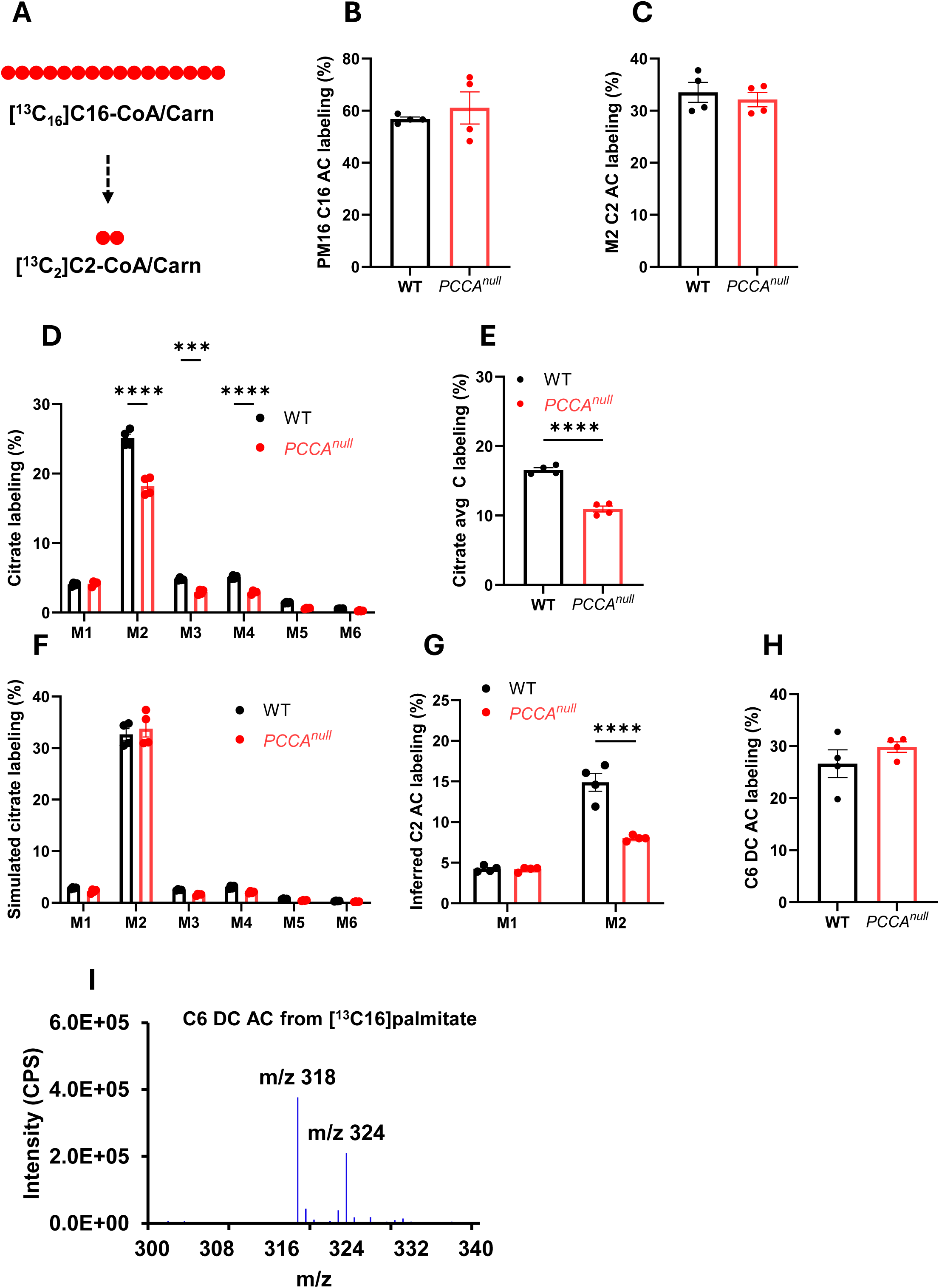
*PCCA* knockout reduces mitochondrial fatty acid oxidation. (A) Schematic of [¹³C₁₆]palmitate metabolism leading to M2 acetyl-CoA/acetylcarnitine. (B–C) Labeling of M16 palmitate and M2 acetylcarnitine in WT and *PCCA^null^*-HepG2 cells following incubation with 0.4 mM [¹³C₁₆]palmitate for 4 hours. (D) Citrate isotopomer distributions in WT and *PCCA^null^*-HepG2 cells under the same conditions. (E) Average carbon (Avg C) labeling of citrate. (F) Simulated citrate labeling based on Equations 1–6 in the Methods. (G) Estimated mitochondrial acetylcarnitine labeling based on Equations 7–9 in the Methods. (H–I) Labeling and mass spectrum of C6 dicarboxylylcarnitine derived from [¹³C₁₆]palmitate. Data are presented as mean ± SEM (n = 4). *, ***, and **** indicate p < 0.05, p < 0.005, and p < 0.0001, respectively.

A marked reduction in pyruvate carboxylation led to a substantial decrease in overall labeling of malate, aspartate, and fumarate, but not succinate (Fig. 3B and Supplementary Figs. 4B, 4D–4F). The labeling pattern of succinate (Supplementary Fig. 4E) more closely resembled that of citrate (Fig. 2F), further supporting that the elevated M3 labeling of malate, aspartate, and fumarate originates from pyruvate carboxylation. These data also suggest that the reverse TCA cycle flux from oxaloacetate (OAA) proceeds only to fumarate (Supplementary Fig. 4G).

Quantitative flux analysis based on citrate isotopomer distributions demonstrated an approximately 82% reduction in the PC/PDH flux ratio in *PCCA^null^*-HepG2 cells (Fig. 3C), indicating a pronounced metabolic shift from anaplerotic to oxidative utilization of pyruvate.

### Reduced pyruvate carboxylation impairs fatty acid synthesis and gluconeogenesis

The high anaplerosis supports anabolic synthesis in the liver cells. The reduction of pyruvate anaplerosis in *PCCA^null^*-HepG2 could affect fatty acid synthesis and gluconeogenesis in HepG2 cells (Fig. 3D). To test these potential consequences *in vivo*, we examined de novo lipogenesis (DNL) in a PA mouse model, *Pcca⁻/⁻* (A138T), and compared it with WT mice (Fig. 3E). Livers from *Pcca⁻/⁻* (A138T) mice showed a significant reduction in DNL relative to WT controls (Fig. 3F).

The abundance of M2-labeled phosphoenolpyruvate (PEP) reflects gluconeogenic activity in the [¹³C₆]glucose tracing experiment. In *PCCA^null^*-HepG2 cells, M2 PEP labeling was significantly reduced compared with WT cells (Supplementary Fig. 3B), indicating impaired gluconeogenesis. This defect was further supported by *in vivo* studies in *Pcca* mutant mice ^34^, suggesting that reduced pyruvate carboxylation is a conserved feature of PCC deficiency. Mechanistically, decreased PC flux may result from reduced mitochondrial acetyl-CoA levels, an allosteric activator of PC, or from diminished ATP availability due to mitochondrial dysfunction.

### *PCCA* deficiency reduces mitochondrial fatty acid oxidation

To evaluate fatty acid metabolism, we performed [¹³C₁₆]palmitate tracing in WT and *PCCA^null^*-HepG2 cells (Fig. 4A and Supplementary Fig. 5A). Labeling of palmitoylcarnitine and intermediate β-oxidation products did not differ significantly between groups, indicating comparable substrate uptake and initial processing (Figs. 4B–4C and Supplementary Figs. 5B–5G). However, incorporation of palmitate-derived carbon into citrate, reflected by both M2 citrate and total citrate labeling, was significantly reduced in *PCCA^null^*-HepG2 cells (Figs. 4D–4E), suggesting impaired entry of fatty acid–derived acetyl-CoA into the TCA cycle. This reduction in [¹³C₁₆]palmitate metabolism was also evident across other TCA cycle intermediates (Supplementary Figs. 6A–6I). Notably, this decrease in TCA cycle labeling from [¹³C₁₆]palmitate in *PCCA^null^*-HepG2 cells was not mirrored by M2 acetylcarnitine labeling (Fig. 2C).

**Figure 5.**
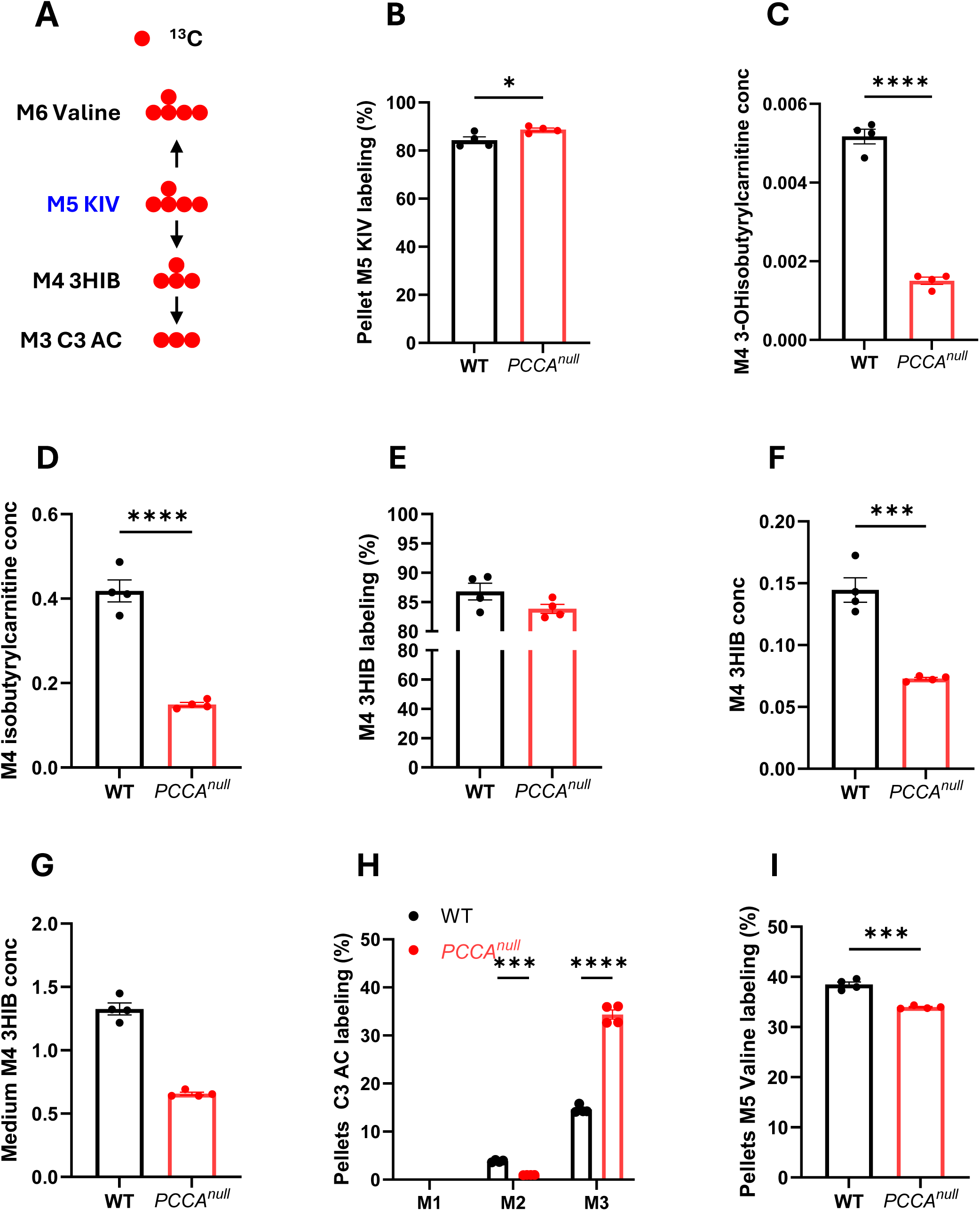
Reduced catabolism of branched-chain keto acids in *PCCA^null^*-HepG2 cells. (A) Schematic of [^13^C_5_]ketoisovalerate (KIV) metabolism. (B) M+5 labeling of KIV in WT and *PCCA^null^*-HepG2 cells following incubation with 0.5 mM [^13^C_5_]KIV for 4 h. (C–D) Levels of M+4 3-hydroxyisobutyrylcarnitine and M+4 isobutyrylcarnitine in WT and *PCCA^null^*-HepG2 cells. (E–G) M+4 3-hydroxyisobutyrate (3-HIB) labeling and abundance in cells and culture medium. (H–I) Propionylcarnitine labeling and M+5 valine labeling in WT and *PCCA^null^*-HepG2 cells after incubation with 0.5 mM [^13^C_5_]KIV for 4 h. Data are shown as mean ± SEM (n = 4). *P < 0.05, ***P < 0.005, ****P < 0.0001.

To resolve this discrepancy, we simulated citrate labeling based on malate and acetylcarnitine isotopomer distributions. The calculation framework is described in Equations 1–6 in the Methods. The simulated M2 citrate showed no difference between groups (Fig. 4D), indicating that total acetylcarnitine labeling does not accurately reflect mitochondrial acetyl-CoA labeling. Using the labeling patterns of malate and citrate, we then estimated mitochondrial acetyl-CoA/acetylcarnitine enrichment (see Equations 7–9 in the Methods). This analysis revealed that mitochondrial M2 acetyl-CoA/acetylcarnitine derived from palmitate was significantly reduced in *PCCA^null^*-HepG2 cells (Fig. 4G).

These findings suggest that a substantial fraction of acetylcarnitine originates from non-mitochondrial sources, most likely peroxisomal fatty acid oxidation. Consistent with this interpretation, we detected labeled dicarboxylic acylcarnitines (C6 dicarboxylylcarnitine; Figs. 4H–4I), a hallmark of peroxisomal metabolism^35–37^. This metabolite was predominantly labeled as M6 in the presence of [¹³C₁₆]palmitate and was absent in unlabeled conditions (Fig. 4I and Supplementary Fig. 5H), confirming its tracer-derived origin.

Together, these results demonstrate that *PCCA* deficiency reduces mitochondrial fatty acid oxidation and limits the contribution of fatty acid–derived acetyl-CoA to the TCA cycle in HepG2 cells.

### Inhibition of branched-chain amino acid catabolism in *PCCA^null^*-HepG2 cells

Propionate is a major resource of propionyl-CoA and its metabolism was attenuated in PA mouse model and hiPSC-CMs ^17,25^. Propiongenic amino acids also contribute to propionyl-CoA production. To assess the impact of PCC deficiency on propiogenic amino acid metabolism, we performed isotope tracing using [¹³C₅]2-ketoisovalerate ([¹³C₅]KIV), a keto acid of branched-chain amino acid (BCAA) catabolism (Fig. 5A). [¹³C₅]KIV was selected over [¹³C]-labeled valine because BCAA metabolism is coordinated between tissues: transamination of BCAAs primarily occurs in muscle, where branched-chain aminotransferase (BCAT) activity is high, generating branched-chain keto acids (BCKAs) that are subsequently transported to the liver for further oxidation, where BCAT activity is relatively low but branched-chain α-keto acid dehydrogenase (BCKDH) activity is high ^38–40^.

While intracellular and extracellular levels of labeled KIV were similar between WT and *PCCA^null^*-HepG2 cells (Fig. 5B and Supplementary Fig. 7A), downstream metabolites, including 3-hydroxyisobutyrylcarnitine, isobutyrylcarnitine, 3-hydroxyisobutyrate (3HIB) were significantly reduced in *PCCA^null^*-HepG2 cells and cultured media (Figs. 5C–5G), indicating impaired BCAA catabolic flux likely due to the feedback by the accumulated propionyl-CoA. Consistent with this, both labeling and accumulation of propionylcarnitine were increased in *PCCA^null^*-HepG2 cells, reflecting a metabolic block downstream of propionyl-CoA. The reduced catabolism of BCAA can also be reflected by the reduction of unlabeled intermediates (Supplementary Figs 7B-7D). In addition, reverse transamination of KIV to valine was reduced as evidenced by the reduced M5 valine labeling (Fig. 5I) and levels (Supplementary 7G), suggesting decreased BCAT activity. Together, these findings indicate an overall suppression of BCAA metabolism, potentially mediated by inhibition of BCKDH activity, which requires further validation.

Tracing with ¹³C-labeled threonine (Fig. 6A) revealed substantial labeling of intracellular threonine but minimal incorporation into propionylcarnitine and TCA cycle intermediates (Figs. 6B–6D and Supplementary Figs. 8A-8D), indicating that threonine contributes minimally to propionyl-CoA production in this system. These results, together with propionate tracing, support the conclusion that propionate is the predominant source of propionyl-CoA in HepG2 cells.

**Figure 6.**
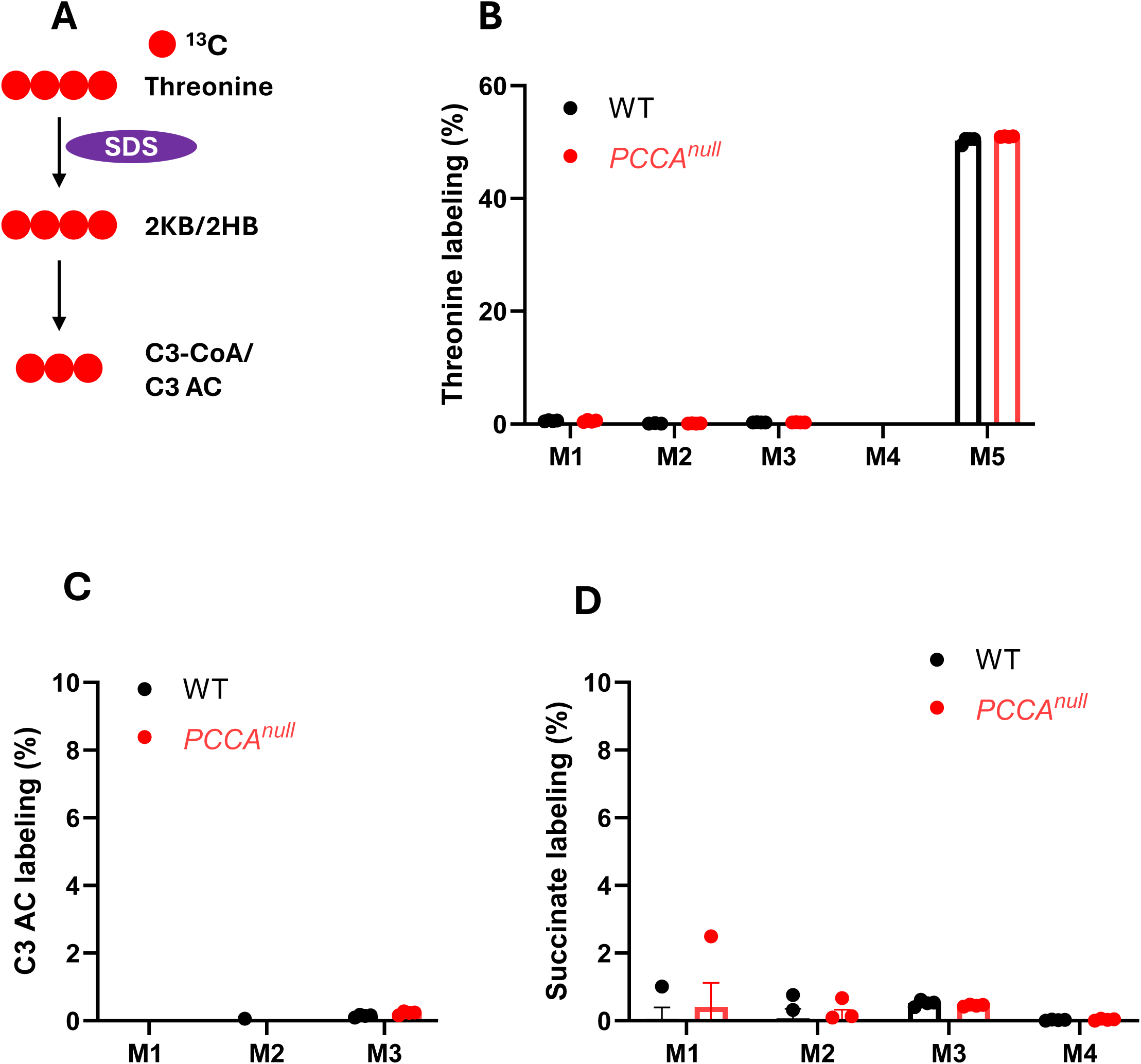
Minimal metabolism of threonine in HepG2 cells. (A) Schematic of [^13^C4]threonine metabolism. (B) Threonine isotopologue labeling in WT and *PCCA^null^*-HepG2 cells following treatment with 0.5 mM [^13^C_4_]threonine for 4 h. (C–D) Stable isotopologue labeling of propionylcarnitine (C3-carnitine) and succinate. Data are shown as mean ± SEM (n = 4).

## Discussions

Patients with PA develop multiple complications even under strict dietary restriction ^8^. The mechanisms underlying these complications remain poorly understood, due in part to the rarity of the disease and limitations of existing animal and cellular models ^17,41,42^. Animal models often fail to fully recapitulate the PA phenotype observed in patients, while human cell lines or organoids combined with genetic manipulations provide a more direct approach for investigating disease-specific metabolic alterations. In this study, we generated a CRISPR knockout of *PCCA* in HepG2 cells to model PA. This system allows amplification of the PA phenotype, offers a human liver-specific context, and avoids confounding contributions from other organs. However, it is important to recognize the limitations that cultured HepG2 cells lack *in vivo* regulatory cues and are derived from human hepatocellular carcinoma, which may influence metabolic behavior. Animal model of PA remains necessary to confirm the findings from *PCCA^null^*-HepG2 cells.

All expected metabolic changes associated with PA were confirmed in *PCCA^null^*-HepG2 cells, with loss of PCC enzyme activity validated by Western blot and enzymatic assays. Interestingly, M3 methylmalonylcarnitine derived from [^13^C_3_]propionate was detectable, albeit at low levels and significantly lower than in WT cells. This suggests that residual conversion of propionyl-CoA to methylmalonyl-CoA may occur via acyl-CoA carboxylase, which warrants further investigation as a potential therapeutic target. Similar to observations in our perfused hearts and hiPSC-derived cardiomyocytes ^17,43^, *PCCA^null^*-HepG2 cells exhibit relatively increased glucose metabolism compared to fatty acid oxidation when propionyl-CoA accumulates. Unlike the cardiac models, where glucose metabolism is increased from glycolysis ^17,43^, *PCCA^null^*-HepG2 cells do not show significant changes in pyruvate, lactate, or alanine labeling from [^13^C_6_]glucose; instead, elevated M2 acetylcarnitine indicates enhanced entry of glucose-derived carbon into the TCA cycle via PDH. This shift may also reflect reduced mitochondrial fatty acid oxidation, consistent with the Randle cycle ^44^, in which decreased fatty acid oxidation promotes glucose utilization as an alternative energy source.

HepG2 cells normally exhibit high pyruvate carboxylase (PC) flux, which provides major anaplerotic support for gluconeogenesis and fatty acid synthesis^45,46^. In *PCCA^null^*-HepG2 cells, PC flux is reduced by nearly 50%, while PDH flux is increased, resulting in a PC/PDH flux ratio that is only 18% of WT levels. Reduced pyruvate anaplerosis likely limits gluconeogenesis and fatty acid synthesis, consistent with impaired gluconeogenesis observed in *Pcca^-/-^*(A138T) mice ^34^. Previous interpretations attributed decreased gluconeogenesis to limited anaplerosis from propionyl-CoA. However, isotope labeling studies with both [^13^C_6_]glucose and [^13^C_3_]propionate show that glucose is the predominant source of OAA, suggesting that reduced gluconeogenesis arises primarily from impaired PC activity. These findings help explain the vulnerability of PA patients to prolonged fasting ^47^.

PCC deficiency also affects fatty acid metabolism in a complex manner. Reduced pyruvate anaplerosis can limit fatty acid synthesis, as evidenced by decreased DNL, whereas inhibited mitochondrial β-oxidation may prevent lipid breakdown, potentially leading to lipid accumulation. This dual effect may underlie conflicting reports in the literature regarding lipid metabolism in PA. Some clinical studies report lipid accumulation ^48,49^, while *Pcca^-/-^*(A138T) mice show reduced fat mass relative to WT animals ^34^, indicating that metabolic outcomes likely depend on additional factors, including environmental and lifestyle conditions.

The observed impairment in pyruvate carboxylation, along with consequent reductions in gluconeogenesis and fatty acid synthesis, mirrors the metabolic disturbances seen in patients with PA, particularly during severe episodes such as metabolic decompensation. These findings also support the clinical rationale for emergency treatment with high glucose and lipid supplementation to counteract catabolism and restore metabolic balance ^1^.

Understanding the major sources of propionyl-CoA is critical for dietary management of PA. While propiogenic amino acids are estimated to contribute over 50% of propionate ^50^, our previous studies demonstrated that microbiome-derived propionate is a major source of propionyl-CoA ^25^, particularly in PA where circulating propionate is markedly elevated. In HepG2 cells, tracing with [^13^C_3_]propionate, [^13^C_5_]KIV, and [^13^C_4_,^15^N]threonine revealed that propionylcarnitine labeling is predominantly derived from propionate (93%), with minor contributions from KIV (14.5%) and threonine (0.17%). These results suggest that controlling propionate levels may be an effective therapeutic approach in PA.

Overall, our human cellular PA model provides direct evidence of metabolic consequences of PCC deficiency using stable isotope-based flux analysis. Reduced pyruvate carboxylation impairs gluconeogenesis and fatty acid synthesis, while reduced mitochondrial fatty acid oxidation drives compensatory glucose oxidation in a less energy-efficient manner. Propionate emerges as the dominant source of propionyl-CoA in HepG2 cells, highlighting both disease-causing mechanisms and potential targets for therapeutic intervention in PA.

## Methods

### Reagents and chemicals

[^13^C_5_]2-Ketoisovaleric acid, [^15^N, ^13^C_4_]threonine, [^13^C_3_]propionate, D9 Carnitine, [^13^C_6_]glucose, and [^13^C_16_]palmitate were from Cambridge Isotope Laboratories (Tewksbury, MA). All other chemicals were from Sigma (St. Louis, MO) using the highest purity.

### Generation of PCCAnull-HepG2 cells

The human PCCA gene (NM_000282.4) and corresponding protein sequence (NP_000273.2) were analyzed to identify suitable CRISPR-Cas9 target sites. Two guide RNAs were designed: T2 (AGAGTCCACAAGGAACTCCA) and T4 (CAGTAAAAGCTACCTCAACA). The endogenous PCCA locus was targeted by transient transfection with CRISPR-Cas9 ribonucleoprotein complexes, resulting in mutations that reduced expression of the encoded protein. Two isogenic knockout clones (Clone 23 and Clone 45) were isolated by limiting dilution in 96-well plates and validated by Sanger sequencing. Biallelic knockout was confirmed by frameshift mutations in both alleles (+1/+1 or -2/-2). Clone 23 was used in the subsequent experiments.

### HepG2 cells culture and tracing experiments

Approximately 1 × 10⁶ WT and *PCCA^null^*-HepG2 cells were plated in 12-well plates and cultured in DMEM supplemented with X% FBS. Stable isotope tracing experiments were then performed using individual tracers in both WT and *PCCA^null^*-HepG2 cells in each experiment. These tracing experiments included 11 mM [¹³C₆]glucose, 1 mM [¹³C₃]propionate, 0.5 mM [¹³C₄,¹⁵N]threonine, and 0.5 mM [¹³C₅]KIV, respectively. For tracing experiments, cells were incubated in DMEM without FBS to minimize interference from unlabeled substrates present in serum. Tracing was conducted for 4 hours, after which cells were washed with cold PBS and harvested in methanol for metabolomic analysis. Pre- and post-incubation media samples were also collected for analysis.

For D₂O tracing experiments, ∼1 × 10⁶ WT and *PCCA^null^*-HepG2 cells were cultured in DMEM containing 3% FBS and 6% D₂O for 3 days. At the end of the labeling period, cells were washed with cold PBS and harvested in methanol for analysis of de novo fatty acid synthesis. Each experiment was performed with four biological replicates.

### DNL mice experiment

All animal procedures were approved by the Duke University IACUC, and all experiments were conducted in accordance with relevant ethical regulations for animal use. The DNL assay using the D₂O labeling approach has been described previously ^40,51,52^. Briefly, WT and *Pcca^-/-^*(A138T) mice (n = 5 per group) received an intraperitoneal bolus of D₂O (10 µL/g body weight), followed by administration of 6% D₂O in drinking water for 2 days prior to sample collection. Anesthesia was induced with 5% isoflurane, and blood samples were collected from both the inferior vena cava and the portal vein. Blood was centrifuged at 12,000 × g for 5 minutes to isolate plasma. At the same time, tissues were rapidly excised, snap-frozen in liquid nitrogen, and pulverized. All samples were stored at −80 °C until further analysis.

### ^2^H_2_O labeling measurement in plasma

The plasma ^2^H_2_O labeling was assayed according to the published protocol ^40,51–53^. Briefly, 10 µl plasma or standard was mixed with 2 µl of a 10 M NaOH solution and 4 µl of acetone/acetonitrile solution (1/20, volume ratio) in Eppendorf tubes. Gently mix samples and incubate overnight (at least for 10 hours). The acetone was extracted by adding 500 µl chloroform. Chloroform phase was dried by ∼50 mg NaSO_4_ salt. Transfer 100 µl chloroform layer into GC vial with glass insert for GC-MS analysis.

### GC-MS method for ^2^H_2_O labeling measurement

Gas Chromatography-Mass Spectrometry (GC-MS): Acetone was analyzed using an Agilent 5973N-MSD equipped with an Agilent 6890 GC system, and a DB-17MS capillary column (30 m x 0.25 mm x 0.25 um). The mass spectrometer is operated in the electron impact mode (EI; 70 eV). The temperature program was as following: 60°C initial, increase by 20°C/min to 100°C, increase by 50°C/min to 220°C, and hold for 1 min. The sample was injected at a split ratio of 40:1 with a helium flow of 1 ml/min. Acetone eluted at 1.5 min. Selective ion monitoring of mass-to-charge ratios of 58 and 59 was performed using a dwell time of 10 ms/ion.

### Total palmitic acid labeling assay in the liver

The total palmitic acid labeling was assayed based on the published work ^40,51–53^. Briefly 20 mg liver tissue was homogenized in 1 ml KOH/EtOH (EtOH 75%) and incubated at 85 °C for 3 hours. A 200 µl of 1 mM [^2^H_31_]palmitate as internal standard was added into samples after cool down. A 100 µl of sample was acidified by 100 µl of 6 M HCl. Palmitic acid was extracted by 600 µl chloroform. Chloroform layer was completely dried by nitrogen gas and was reacted with 50 µl N-tert-Butyldimethylsilyl-N-methyltrifluoroacetamide (TBDMS) at 70 °C for 30 minutes. The TBDMS-derivatized samples were ready for GC-MS analysis. Gas Chromatography-Mass Spectrometry (GC-MS): Palmitate –TBDMS derivative was analyzed using an Agilent 5973N-MSD equipped with an Agilent 6890 GC system, and a DB-17MS capillary column (30 m x 0.25 mm x 0.25 µm). The mass spectrometer was operated in the electron impact mode (EI; 70 eV). The temperature program was as follows: 100°C initial, increase by 15°C/min to 295°C and hold for 8 min. The sample was injected at a split ratio of 10:1 with a helium flow of 1 ml/min. Palmitate-TBDMS derivative eluted at 9.7 min. Mass scan from 100 to 600 was chosen in the method. The m/z at 313, 314, and 344 were extracted for M0, M1, and M31 palmitate quantification.

### DNL calculation

All the stable isotope labeling was corrected from the natural stable isotope distribution^54^. The newly synthesized total palmitic acid was calculated as following ^51–53^: %newly synthesized palmitic acid labeling = total palmitic acid labeling /(plasma ^2^H_2_O labeling × 22)×100

### PCCA expression by Western Blot

The Western Blot was modified from our previous work ^21^. Approximately 1 million HepG2 cells from WT and *PCCA^null^* were lysed in 500 µl RIPA buffer with protease inhibitors. Centrifuge for 5 minutes at 4 °C to clarify. Take lysate containing 20 µg protein and mix with loading buffer (Bio-RAD, 4× Laemmli Sample Buffer with 10% 2-mercapethanol). Heat lysate at 90 °C for 5 minutes and load the above lysate samples, protein marker (5 µl), and PCCA lysate (PCCA standard, 2 µl of 20 µg/50 µl in loading buffer) to Precast Gel (Mini-PROTEAN TGX stain- Free). Run gel for 30 minutes at constant voltage of 200 V. Transfer proteins from gel to PVDF membrane for 10 minutes. The membrane was washed with 1X PBS buffer for 5 minutes and blocked for 1 h with 5% milk buffer in TBS. And then expose the membrane to primary antibody anti-PCCA (Dilution 1:3000, Protein Tech.) at 4 °C on a shaker overnight. Wash membrane with 1XTBST (4 times for 5 minutes). Membrane was incubated with secondary antibody (the diluted IRDye Goat anti-Rabbit, 1:10000) for 1 hour and was protected from light. Then wash extensively (4 times for 5 minutes) and rinse in 1XPBS for 5 minutes, followed with scanning on an Odyssey Imaging System. Beta-tubulin was used as a loading control. Empty the buffer and add 10 ml stripping buffer (1x re-blot plus mild solution). Shake at room temperature for 30 minutes. Wash the membrane with 1X TBST for 4 times, then incubate with Tubulin antibody (Sigma, made in mouse) 1:20000 at 4 °C for overnight. Wash the membrane with 1XTBST (4 times for 5 minutes) and then incubate with the secondary antibody (the diluted IRDye 800CW Goat anti-Mouse, 1:10000) for 1 h after being protected from light. The membrane was washed extensively (4 times for 5 minutes) and was rinsed in 1XPBS for 5 minutes. The membrane was scanned on an Odyssey imaging system.

### GC-MS for metabolite profile

We profiled the metabolic changes in cell lystate using our previously published GC-MS method ^17,21,25,34,43,55^. Briefly, a lysate sample was spiked with 0.2 nmol of norvaline and 0.4 nmol [^2^H_9_]L-carnitine or mixed stable isotope labeled metabolites as internal standards and 1000 μl methanol was added and homogenized by sonication for 1 minute. Centrifuged for 20 minutes. The upper phase, approximately 300 µl in volume, was transferred to a fresh Eppendorf vial and subsequently evaporated using nitrogen gas. The resulting dried residues underwent sequential derivatization with methoxylamine hydrochloride and N-tert-butyldimethylsilyl-N-methyltrifluoroacetamide (TBDMS). Specifically, 40 μl of methoxylamine hydrochloride (2% (w/v) in pyridine) was added to the dried residues, followed by incubation for 90 minutes at 40°C. Subsequently, 20 μl of TBDMS with 1% tert-butylchlorodimethylsilane was added, and the mixture was incubated for an additional 30 minutes at 80°C. The derivatized samples were then centrifuged for 10 minutes at 12,000 × g, and the supernatants were transferred to GC vials for further analysis. For GC/MS analysis, we employed an Agilent 7890B GC system with an Agilent 5977A Mass Spectrometer, following the methodology described in our previous work 49, 58, 59. Specifically, 1 µl of the derivatized sample was injected into the GC column. The GC temperature gradient began at 80°C for 2 minutes, increased at a rate of 7°C per minute to 280°C, and was maintained at 280°C until the 40-minute run time was completed. The ionization was conducted via electron impact (EI) at 70 eV, with Helium flow at 1 ml min-1. Temperatures of the source, the MS quadrupole, the interface, and the inlet were maintained at 230°C, 150°C, 280°C, and 250°C, respectively. Mass spectra (m/z) in the range of 50 to 700 were recorded in mass scan mode.

### LC-MS/MS for acylcarnitine profile

Cell lysate acylcarnitines were methylated and profiled using a modified LC-MS/MS method ^17,21,25,34,43,55^. The lysate sample extracts (300 µl) from the previous sample preparation were completely dried using nitrogen gas. The dried residues were then methylated with a 3 M HCl methanol solution (100 µl) at 50°C for 25 minutes. After methylation, the samples were once again dried completely using nitrogen gas and then reconstituted in 20 µl of methanol and 60 µl of water. The derivatized samples were subsequently analyzed using an LC-QTRAP 6500+-MS/MS (Sciex, Concord, Ontario). A gradient HPLC method with two mobile phases (mobile phase A was 98% water with 2% acetonitrile and 0.1% formic acid and mobile phase B was 98% acetonitrile with 2% H_2_O and 0.1% formic acid) was adopted to run with an Agilent Pursuit XRs 5 C18 column (150 × 2.0 mm). The gradient started with 0% B within the first 2 minutes and then increased to 80% at 13 minutes. The column was washed out by 90% B for 4 minutes and equilibrated with initial condition (2% B) for 5 minutes before next injection. The flow rate was 0.4 ml minute-1 and the column oven was set at room temperature. The injection volume was 2 µl. The parameters for Sciex QTRAP 6500+ mass spectrometry were optimized as follows: DP: 33 V, EP 10 V, CXP: 10 V, source temperature: 680 °C, gas 1: 65, gas 2: 65, curtain gas: 35, CAD: 10, and ion spray voltage: 5500 V. The Q1 of all the methylated acylcarnitines was scanned from m/z 218 to m/z 444 with the same fragment (Q3) at m/z 99. L-carnitine had the ion transition of Q1 (m/z 176) and Q3 (m/z 85 or m/z 117). [^2^H_9_]L-carnitine has the shifted Q1 at m/z 179 or m/z 185 with the same Q3 at m/z 85 or m/z 117.

### PCC activity assay

The approach to measure PCC activity was adopted from published work ^34^. Ten microliters of cell lysate (∼4 mg protein/ml) were used, and 100 μl of an enzyme reaction mixture was added. This mixture consisted of 100 mM Tris-HCl (pH 7.5), 5 mM MgCl_2_, 1 mM DTT, 10 mM KCl, 40 mM NaHCO_3_, 1 mM Biotin, and 6 mM ATP. The reaction was initiated by adding 10 μl of 11.8 mM propionyl-CoA and carried out at 37°C for 30 minutes. Subsequently, 10 μl of the reaction mixture was combined with 10 μl of 0.01 mM D9 pentanoyl-CoA as an internal standard, followed by mixing with 50 μl of 200 mM formic acid to terminate the reaction. The mixture was then centrifuged at 13,000 × rpm for 10 minutes. The supernatants were analyzed using LC-QTRAP 6500^+^-MS/MS (Sciex, Concord, Ontario). Ion chromatograms of propionyl-CoA, methylmalonyl-CoA, and D9 pentanoyl-CoA were extracted and quantified based on m/z values of 824/317, 868/361, and 861/354, respectively.

### Calculations and statistics

Measured mass isotopologues distributions expressed as mol percent were corrected for natural enrichment^54^. M0, M1, M2, …, Mn represents the isotopologues of molecule with n heavy atoms. Statistical differences were analyzed by Prism software. Student’s T test was employed when two groups of data were compared. Two-way ANOVA analysis followed by Post Hoc (Turkey) test was conducted for data from more than two groups.

#### Inference of citrate and mitochondrial acetylcarnitine (AC) labeling from malate (MA) labeling

Citrate (Cit) labeling was simulated based on the measured labeling of malate and acetylcarnitine using the following relationships:

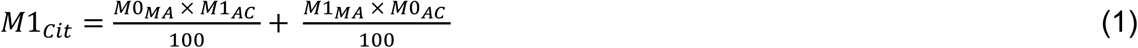

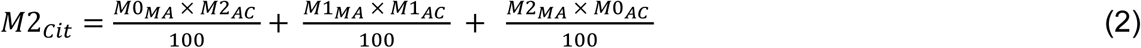

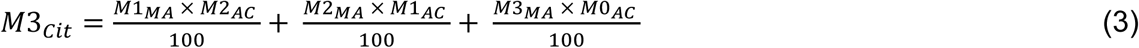

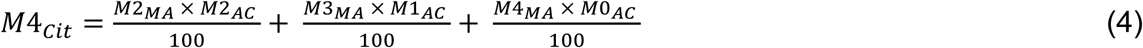

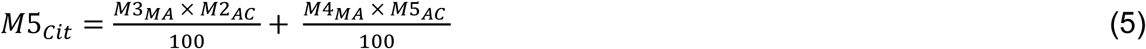

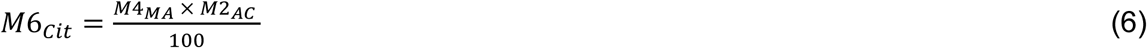

M1_Cit_, M2_Cit_, M3Cit, M4Cit, M5Cit, and M6Cit represent the labeling of citrate isotopomers. M0MA, M1MA, M2MA, M3MA, and M4MA represent the labeling of malate isotopomers. M0AC, M1AC, and M2 AC represent the labeling of acetylcarnitine isotopomers.

#### Mitochondrial acetylcarnitine labeling calculation

To estimate the mitochondrial acetylcarnitine labeling, M0, M1, and M2 isotopomers of acetylcarnitine were inferred by solving the following system of three equations:

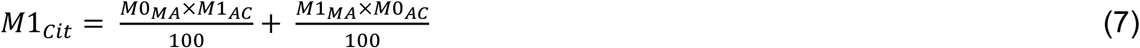

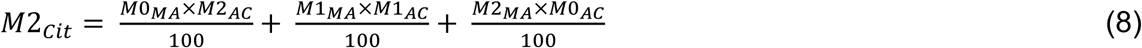

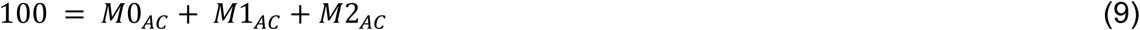

All calculations in Equations 1–9 are based on the assumption that labeling between oxaloacetate (OAA) and malate, as well as between acetyl-CoA and acetylcarnitine, is fully equilibrated.

## Acknowledgement

This work is supported by NIH R21 TR005163 (GFZ and XC), WW Smith Foundation H2504 (XC) and Propionic Acidemia Foundation (GFZ).

## Author Contributions

F.L. C.P., W.H., Z.X., and P.T. performed all experiments. T.K., X.C., and G.F.Z. participated data analysis, manuscript writing, and editing. All authors read and approved the manuscript in its final form.

## Data availability

All data supporting the findings of this study are included in the article and its Supplementary information. Numerical source data for the graphs in the manuscript are available in Supplementary Data. Other data that support the findings of this study are available from the corresponding author upon request.

## Competing Interests

The authors declare no competing interests.

## Abbreviations

PA: propionic acidemia
PCC: propionyl-CoA carboxylase
TCA cycle: tricarboxylic acid cycle
hiPSC: human induced pluripotent stem cells
PDH: pyruvate dehydrogenase
PC: pyruvate carboxylase
WT: wild type
DNL: de novo lipogenesis
PEP: phosphoenolpyruvate
OAA: oxaloacetate
BCAA: branched-chain amino acid
BCKA: branched-chain keto acids
KIV: 2-ketoisovalerate
BCKDH: branched-chain α-keto acid dehydrogenase
BCAT: branched-chain amino acid aminotransferase
3HIB: 3-hydroxyisobutyrate

---

**Supplementary Figure 1.**
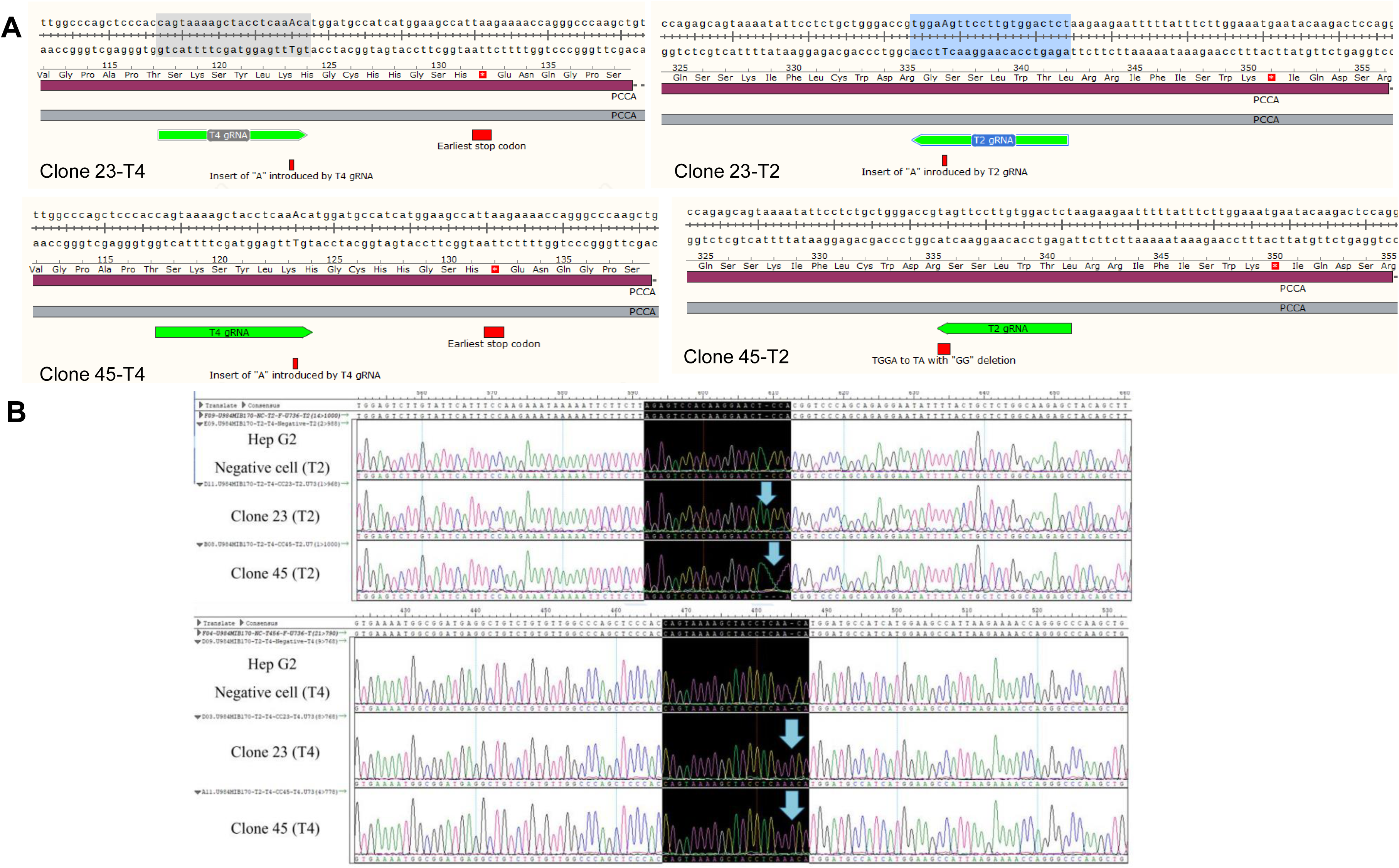
Biallelic knockout of the *PCCA* gene in HepG2 cells. (A) Schematic representation of the guide RNAs and the resulting mutations in two isogenic knockout clones. (B) Sanger sequencing traces of the two isogenic knockout clones compared with parental HepG2 cells.

**Supplementary Figure 2.**
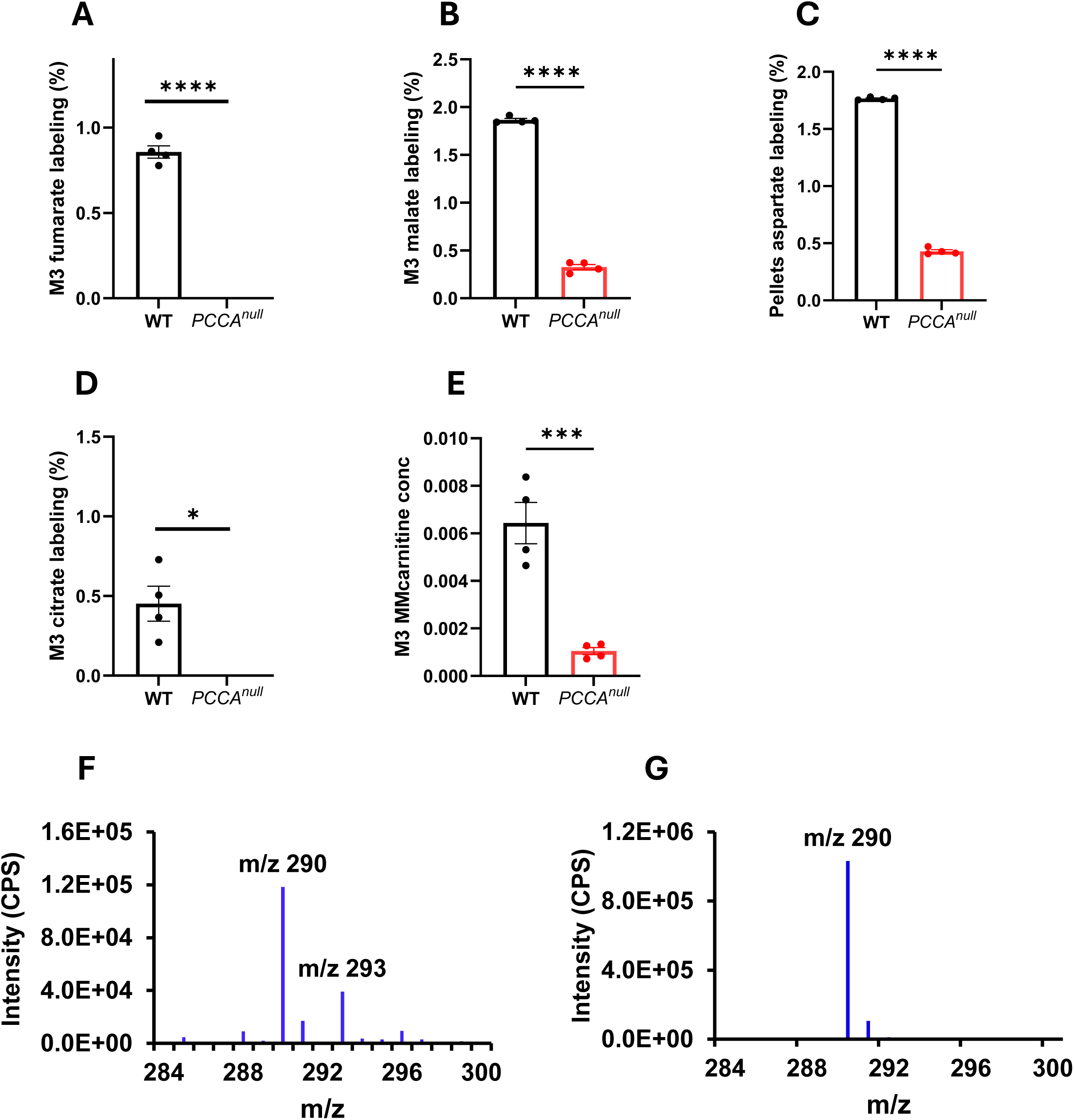
Reduced propionyl-CoA metabolism in *PCCA^null^*-HepG2 cells. (A–D) M3 isotopomer labeling of fumarate, malate, aspartate, and citrate in wild-type (WT) and *PCCA^null^*-HepG2 cells following incubation with 1 mM [¹³C₃]propionate for 4 hours. (E) Abundance of M3 methylmalonylcarnitine (MM-carnitine) in WT and *PCCA^null^*-HepG2 cells after 4-hour incubation with 1 mM [¹³C₃]propionate. (F–G) Mass spectra of methylmalonylcarnitine in HepG2 cells cultured with or without [¹³C₃]propionate. Data are presented as mean ± SEM (n = 4). *, ***, and **** indicate p < 0.05, p < 0.005, and p < 0.0001, respectively.

**Supplementary Figure 3.**
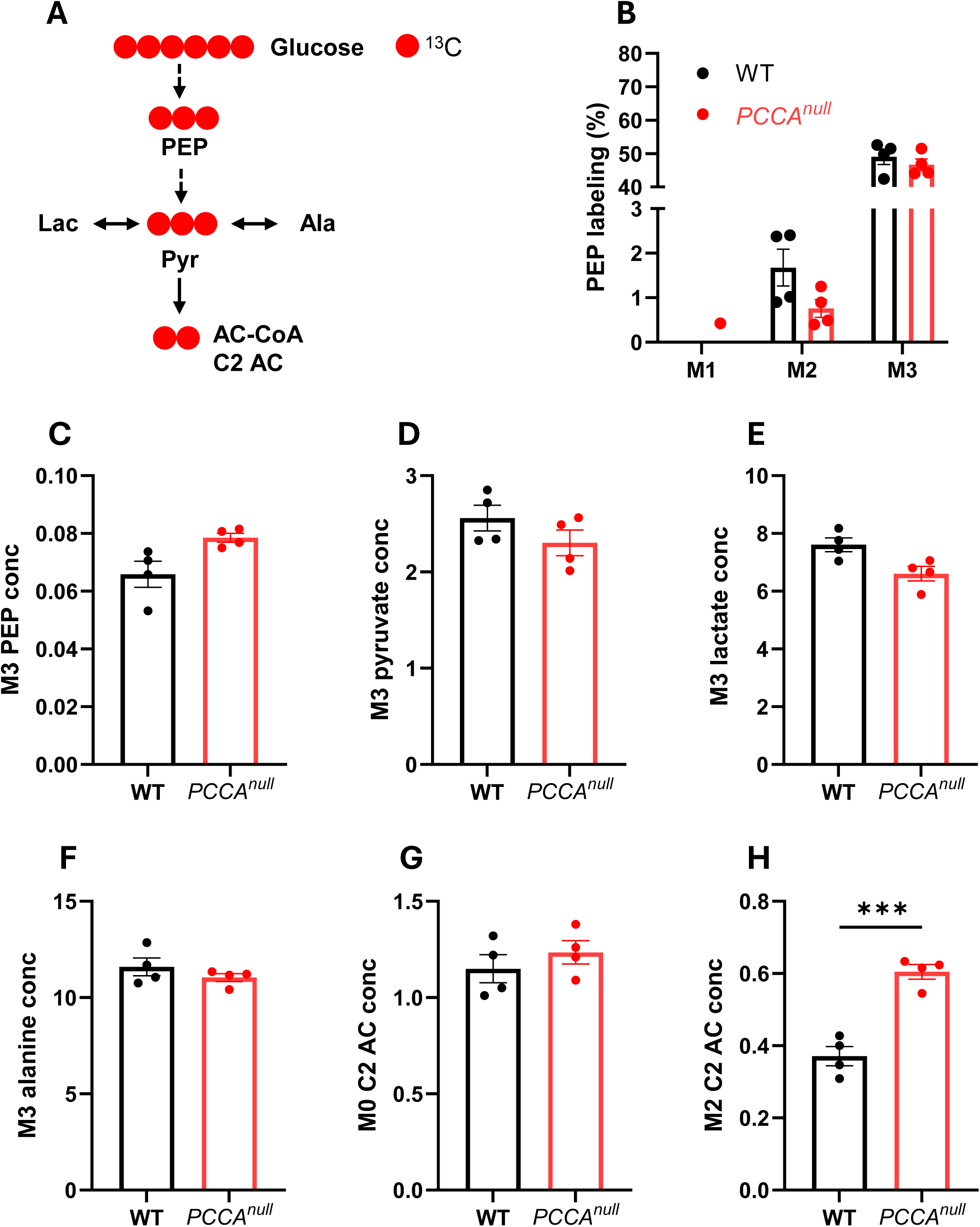
*PCCA* knockout enhances glucose metabolism through PDH. (A) Schematic of glycolysis and the generation of acetyl-CoA/acetylcarnitine from [¹³C₆]glucose. (B) Isotopomer distribution of phosphoenolpyruvate (PEP). (C–H) Levels of M3 PEP, M3 pyruvate, M3 lactate, M3 alanine, M0 acetylcarnitine, and M2 acetylcarnitine in wild-type (WT) and *PCCA^null^*-HepG2 cells. Data are presented as mean ± SEM (n = 4). **** indicates p < 0.0001.

**Supplementary Figure 4.**
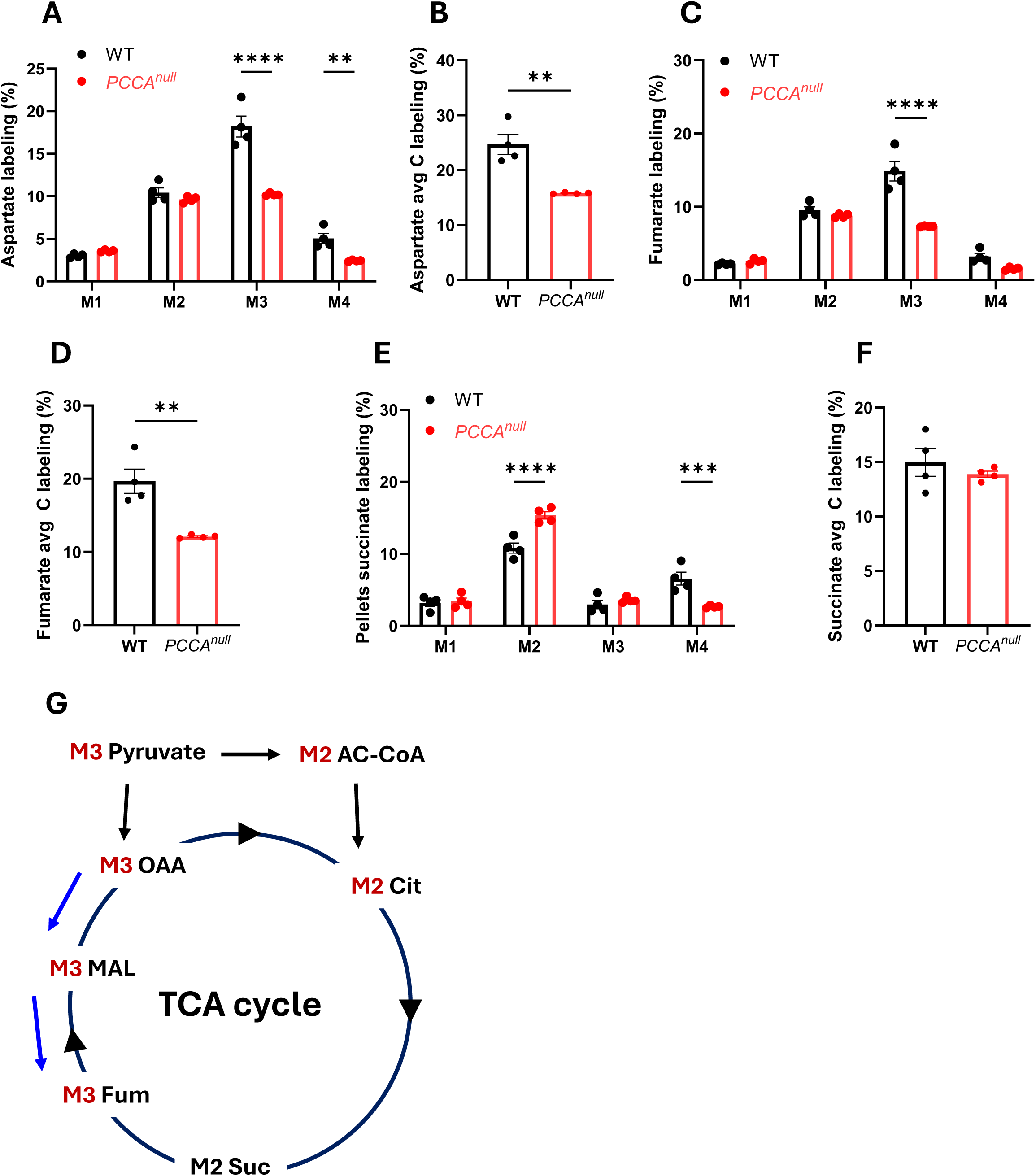
*PCCA* knockout reduces pyruvate anaplerosis in HepG2 cells. (A–F) Mass isotopomer distributions and average carbon (avg C) labeling of aspartate, fumarate, and succinate in wild-type (WT) and *PCCA^null^*-HepG2 cells following incubation with 11 mM [¹³C₆]glucose for 4 hours. (G) Schematic illustrating the predominant isotopomers of TCA cycle intermediates derived from [¹³C₆]glucose via pyruvate carboxylation and pyruvate dehydrogenase flux. Data are presented as mean ± SEM (n = 4). ** and **** indicate p < 0.01 and p < 0.0001, respectively.

**Supplementary Figure 5.**
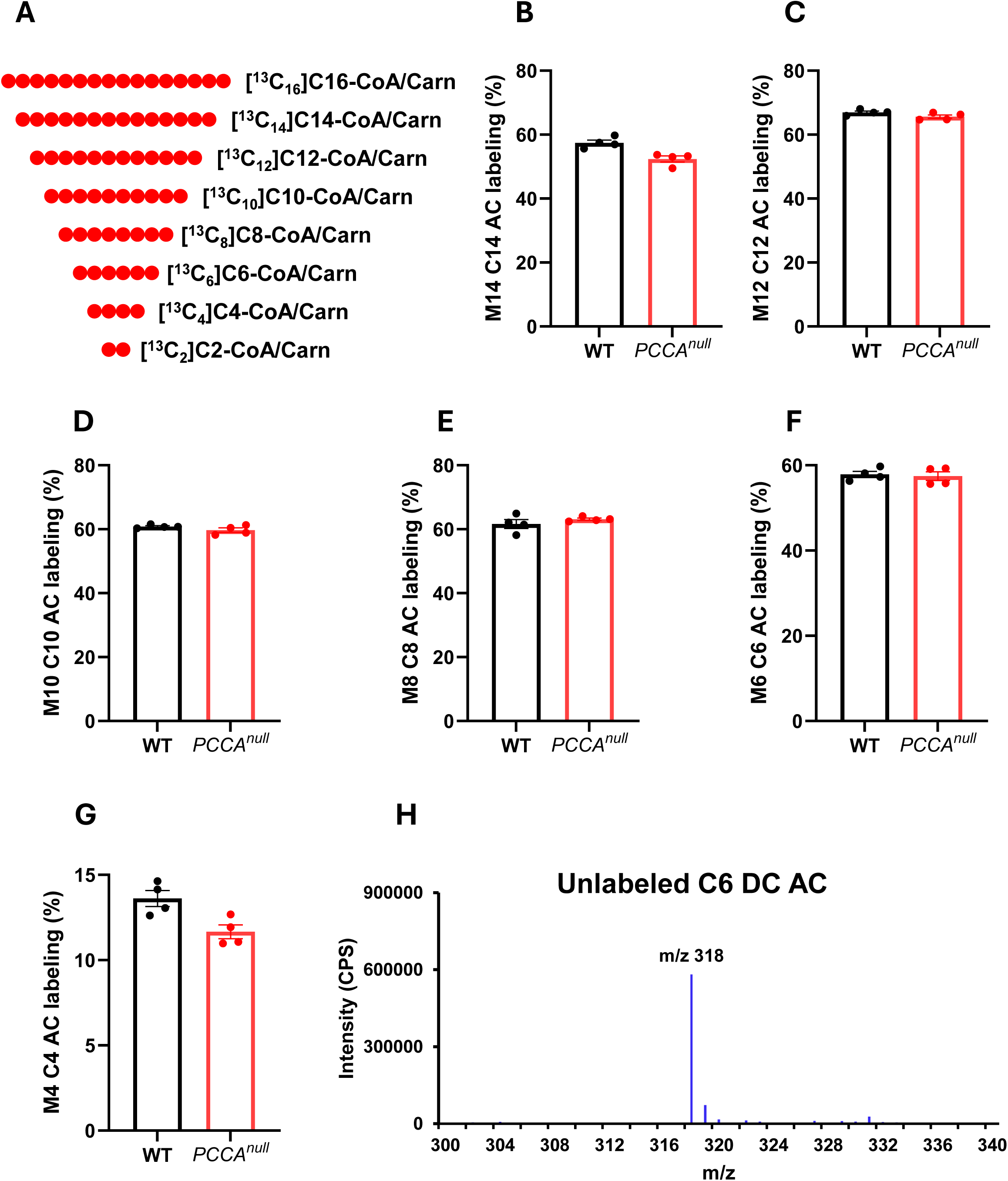
Unchanged acylcarnitine labeling derived from [¹³C₁₆]palmitate in *PCCA^null^*-HepG2 cells. (A) Schematic of acylcarnitine intermediates generated along the fatty acid β-oxidation pathway. (B–G) Labeling of M14 C14 acylcarnitine (C14 AC), M12 C12 AC, M10 C10 AC, M8 C8 AC, M6 C6 AC, and M4 C4 AC in wild-type (WT) and *PCCA^null^*-HepG2 cells following incubation with 0.4 mM [¹³C₁₆]palmitate for 4 hours. (H) Mass spectrum of unlabeled C6 dicarboxylylcarnitine (C6 DC AC). Data are presented as mean ± SEM (n = 4).

**Supplementary Figure 6.**
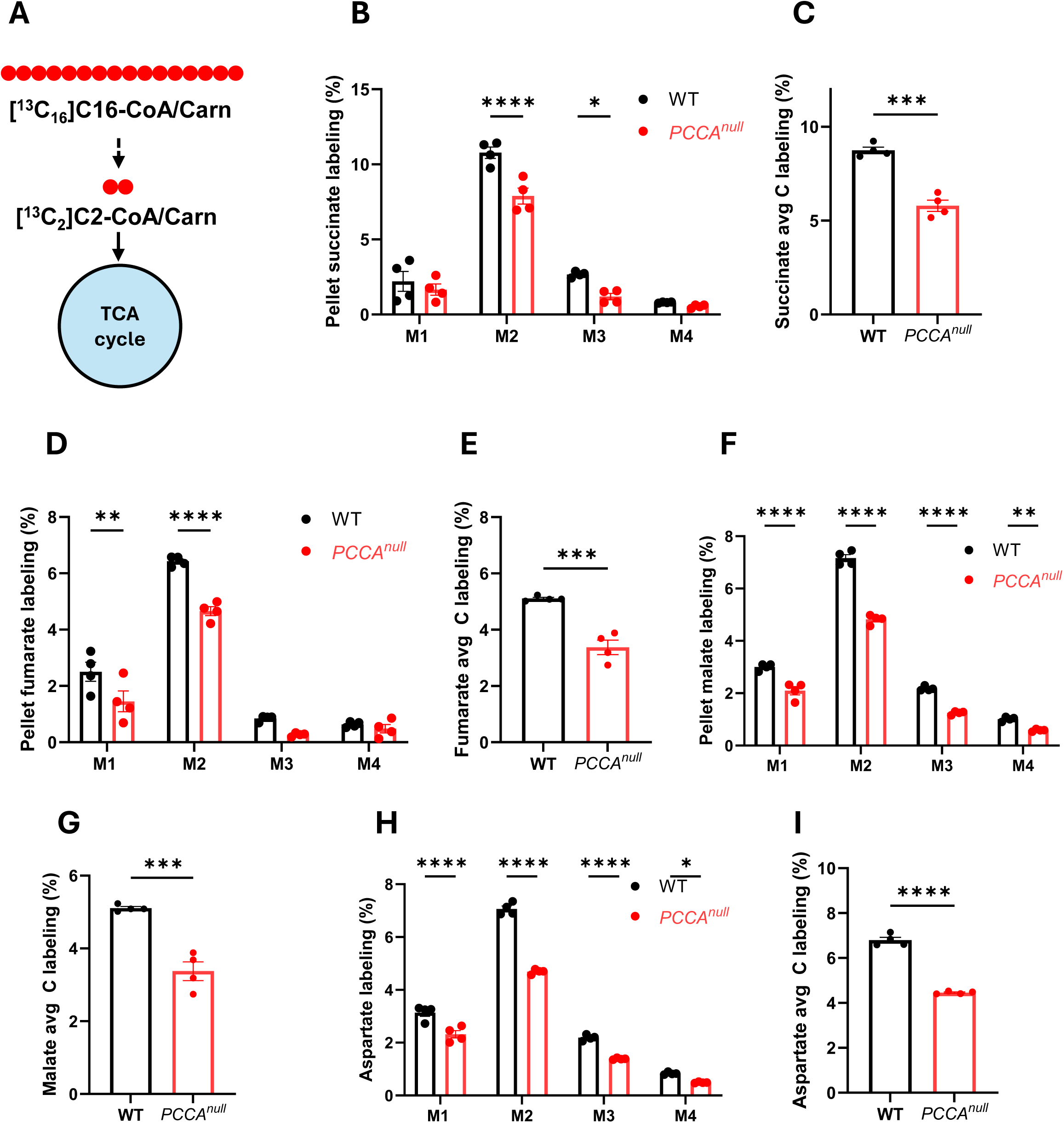
Reduced mitochondrial fatty acid oxidation in *PCCA^null^*- HepG2 cells. (A) Schematic of [¹³C₁₆]palmitate metabolism entering the TCA cycle. (B–I) Stable isotopomer distributions and average carbon (Avg C) labeling of succinate, fumarate, malate, and aspartate in wild-type (WT) and *PCCA^null^*-HepG2 cells following incubation with 0.4 mM [¹³C₁₆]palmitate for 4 hours. Data are presented as mean ± SEM (n = 4). *** and **** indicate p < 0.005 and p < 0.0001, respectively.

**Supplementary Figure 7.**
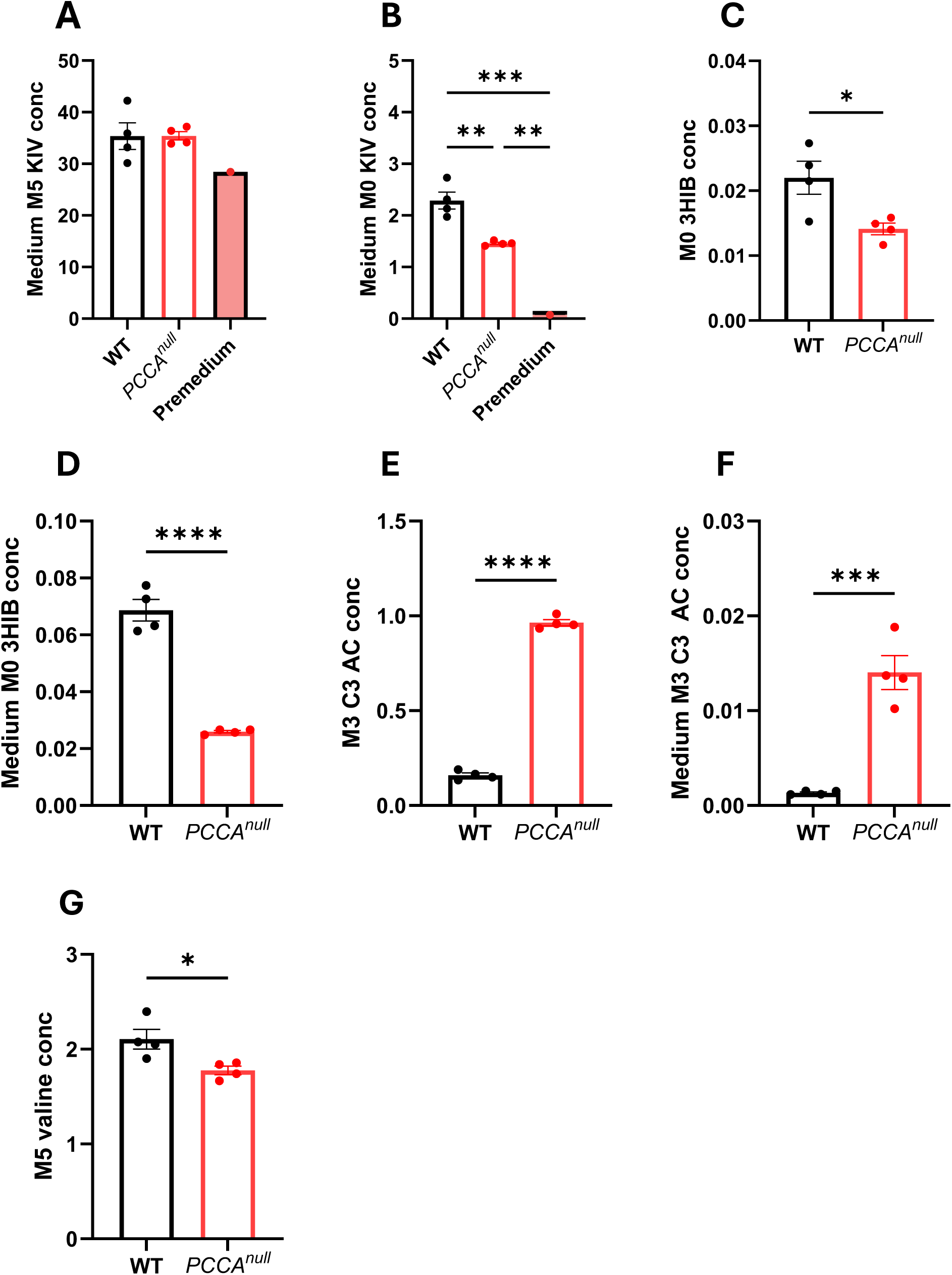
PCCA deficiency reduces branched-chain keto acid (BCKA) catabolism. (A–B) M5 and M0 KIV levels in pre-incubation and conditioned media from WT and *PCCA^null^*-HepG2 cells. (C–F) Unlabeled 3-hydroxyisobutyrate (3-HIB) and M3 propionylcarnitine (C3-carnitine) levels in cell pellets and culture media. (G) M5 valine levels in WT and *PCCA^null^*-HepG2 cells. Data are shown as mean ± SEM (n = 5). *P < 0.05, **P < 0.01, ***P < 0.005, ****P < 0.0001.

**Supplementary Figure 8.**
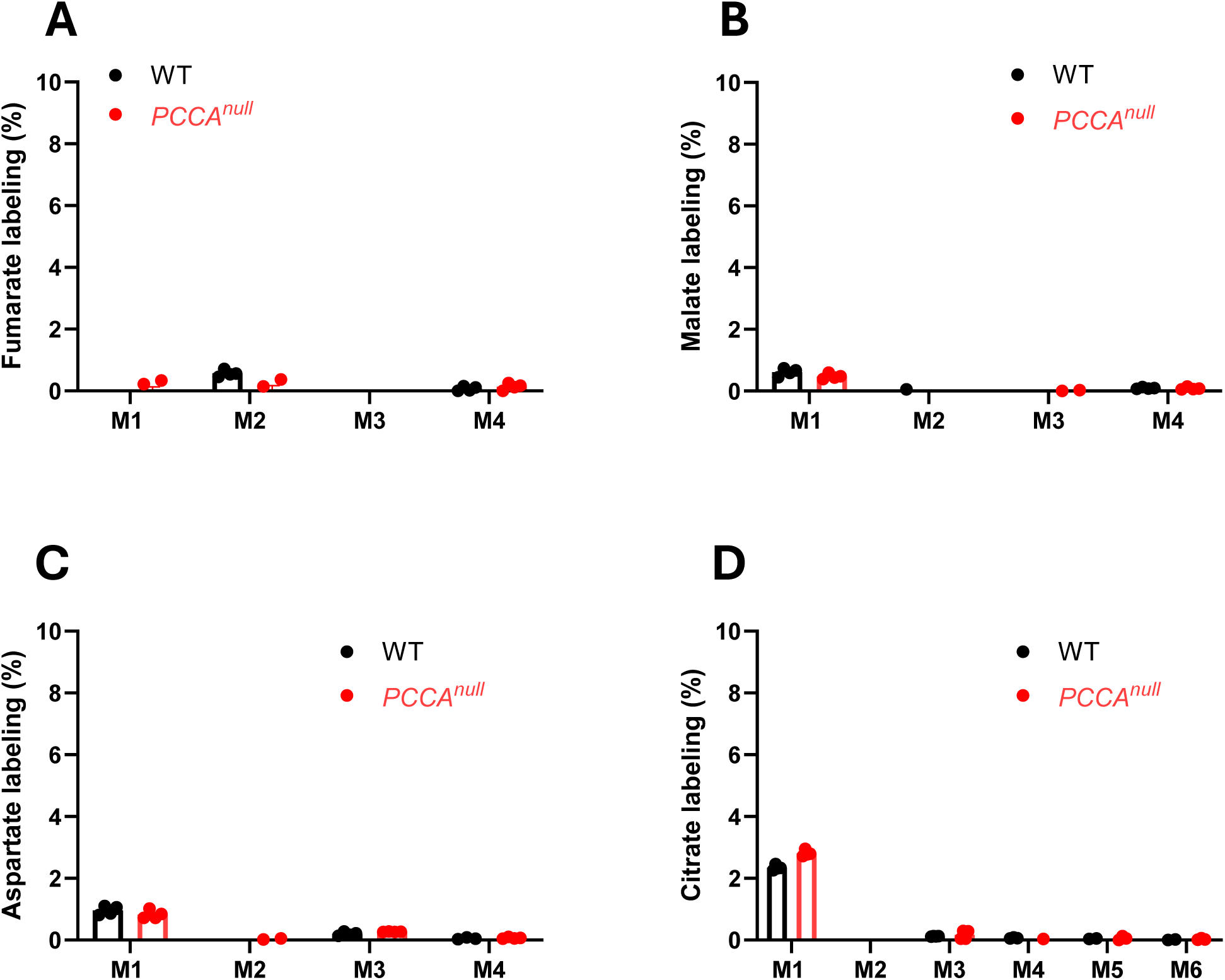
Minimal metabolism of threonine in HepG2 cells. (A–D) Stable isotopologue labeling of fumarate, malate, aspartate, and citrate in WT and *PCCA^null^*-HepG2 cells following treatment with 0.5 mM [^13^C_4_]threonine for 4 h. Data are shown as mean ± SEM (n = 4).

## Notes

### Competing Interest Statement

The authors have declared no competing interest.

